# Perinatal Fentanyl Exposure Drives Enduring Addiction Risk and Central Amygdala Gene Dysregulation

**DOI:** 10.1101/2025.05.20.655167

**Authors:** Courtney P. Wood, Yeji Shin, Maria G. Balaguer, Paola Campo, Selen Dirik, Bryan A. Montoya, Gregory M. R. Cook, Gabrielle M. Palermo, Parsa K. Naghshineh, Alex Morgan, Sara R. M. U. Rahman, Abraham A. Palmer, Francesca Telese, Giordano de Guglielmo

## Abstract

The use of fentanyl and other opioids during pregnancy is a pressing public health issue due to its association with Neonatal Opioid Withdrawal Syndrome (NOWS) and long-term neurobehavioral deficits. Human epidemiologic studies are confounded by both genetic and environmental factors that differ between exposed and unexposed children. We developed a novel rat model of perinatal fentanyl exposure to characterize NOWS symptoms and investigate enduring behavioral and molecular outcomes. Offspring born to fentanyl-exposed dams exhibited reduced survival, lower body weight, spontaneous withdrawal symptoms, and mechanical hypersensitivity. In adolescence, these rats displayed negative affect, while in adulthood, they showed increased fentanyl self-administration, heightened drug-seeking during reinstatement, and elevated corticosterone levels during withdrawal. To explore the molecular underpinnings of these physiological and behavioral outcomes, we conducted RNA-seq in the central amygdala of adult rats, revealing dysregulated pathways related to GPCR signaling, adaptive immune response and neurodevelopmental processes. These transcriptional changes provide insights into the mechanisms driving addiction vulnerability and stress-related behaviors following early fentanyl exposure. Our findings highlight the lasting impact of perinatal opioid exposure in an experimental system that avoids many of the confounds that plague studies in humans, underscoring the need for preclinical models to study NOWS and its long-term consequences. This model offers translational relevance for developing therapeutic strategies to mitigate NOWS and reduce neuropsychiatric risks associated with prenatal opioid exposure.

## Introduction

The rise in opioid use during pregnancy has escalated concerns over Neonatal Opioid Withdrawal Syndrome (NOWS) (Patrick et al., 2012), a condition affecting at least 90 infants each day in the United States. Newborns diagnosed with NOWS experience distressing symptoms including difficulties with sleeping and feeding, muscle twitches, seizures, vomiting, and diarrhea (Ko et al., 2017). Beyond these, long-term neurodevelopmental deficits, such as hyperactivity, impulsivity, aggression, and altered stress reactivity, persist into childhood (Beckwith and Burke, 2015; Hunt et al., 2008; Ornoy, 2003; Ornoy et al., 2001; Wilson et al., 1979). Given the rising rates of opioid use during pregnancy (Haight et al., 2024; Ko et al., 2017; Villapiano et al., 2017), NOWS has emerged as a critical issue. However, studies in humans cannot discriminate between the direct effects of *in utero* opioid exposure and the genetic and environmental differences between exposed and unexposed children, highlighting an urgent need for preclinical models that can accurately mimic perinatal opioid exposure while avoiding the confounding factors that are present in human studies in which *in utero* exposure is not randomly assigned.

*In utero* opioid exposure leads to structural, functional, neurophysiological, and circuit-based disruptions during brain development (Yen and Davis, 2022). Critical structures involved in reward, impulse control, and decision-making have been shown to develop incorrectly in opioid-exposed infants (Lust et al., 2024; Merhar et al., 2021; Rana et al., 2019; Schulson et al., 2014; Yuan et al., 2014). These changes may predispose offspring to addiction-like behaviors (Crews et al., 2007; Dayan et al., 2010; Fareri et al., 2008). While prior studies focused on morphine or oxycodone, the rise of fentanyl, a highly potent synthetic opioid linked to escalating overdose deaths, remains understudied in perinatal contexts (Kuczynska et al., 2018; Warner et al., 2016). To address this gap, we developed a rat model of perinatal fentanyl exposure to examine NOWS symptoms, long-term behavioral outcomes, and molecular changes. We assessed fentanyl self-administration and relapse-like behavior in adulthood, alongside transcriptomic profiling of the central amygdala, a key region in addiction and affect. These findings elucidate mechanisms underlying addiction vulnerability and inform therapeutic strategies for NOWS.

## Materials & Methods

### Animals

Heterogeneous Stock (HS) rats (n=16, 8 males and 8 females) were bred at University of California San Diego in the laboratory of Dr. Abraham A. Palmer (colony identifier: McwiWfsmAap:HS, [RID:RGD_155269102](https://rgd.mcw.edu/rgdweb/report/strain/main.html?id=155269102)). Rats were housed two per cage on a 12 hr/12 hr reversed light/dark cycle in a temperature (20–22°C) and humidity (45–55%) controlled vivarium with ad libitum access to tap water and standard rodent food pellets. Female rats (12-14 weeks old) were surgically implanted with subcutaneous minipumps (see *Drug Preparation and Administration)* and allowed to recover for one week prior to being placed in breeding pairs. Once dams became pregnant, sires were removed from the cage. Offspring were weaned at PND 28, at which time they stopped receiving fentanyl through the dam’s breastmilk. All procedures were conducted in strict adherence to the National Institutes of Health Guide for the Care and Use of Laboratory Animals and were approved by the Institutional Animal Care and Use Committees of UC San Diego (S21145).

### Drug Preparation and Administration

Fentanyl hydrocholoride (Cayman Chemicals, Ann Arbor, MI, USA) was diluted in 0.9% sterile saline and injected into sterile osmotic minipumps (Alzet, Cupertino, CA, USA). Minipumps began releasing fentanyl 24 hours after surgical insertion at a dose of 0.6 mg/kg/day (2.5 µL per hour) for 28 consecutive days. For intravenous self-administration, fentanyl hydrochloride was diluted in 0.9% sterile saline at a dose of 3.2 µg/kg/infusion. To ensure consistent dosing, animals were weighed weekly to adjust the drug solution concentration, rounded to the nearest ten grams.

### Minipump Implantation

Prior to being paired with breeding partners, dams (12-14 weeks old) were anesthetized with vaporized isoflurane (1-5%). Pre-filled osmotic minipumps were aseptically inserted subcutaneously via a horizontal incision on the dorsal surface of the rat. The incision was sutured closed with surgical silk and rats were allowed to recover for one week prior to being introduced to a breeding partner. For 5 days following surgery, flunixin (2.5 mg/kg, s.c.) was administered as analgesic, and cefazolin (330 mg/kg, i.m.) as antibiotic.

### Intravenous Catheterization

Fentanyl- and vehicle-exposed offspring (8-10 weeks old) were anesthetized with vaporized isoflurane (1–5%). Intravenous catheters were aseptically inserted into the right jugular vein using the procedure described previously (Kallupi et al., 2020). Catheters consisted of Micro-Renathane tubing (18 cm, 0.023-inch inner diameter, 0.037-inch outer diameter; Braintree Scientific, Braintree, MA, USA) attached to a 90-degree angle bend guide cannula (Plastics One, Roanoke, VA, USA), embedded in dental acrylic, and anchored with mesh (1 mm thick, 2 cm diameter). Tubing was inserted into the vein following a needle puncture (22 G) and secured with a suture. The guide cannula was punctured through a small incision on the back. The outside part of the cannula was closed off with a plastic seal and metal cover cap, which allowed for sterility and protection of the catheter base. Flunixin (2.5 mg/kg, s.c.) was administered as analgesic, and Cefazolin (330 mg/kg, i.m.) as antibiotic. Rats were allowed one week for recovery prior to any self-administration. They were monitored and flushed daily with heparinized saline (10 U/ml of heparin sodium; American Pharmaceutical Partners, Schaumberg, IL, USA) in 0.9% bacteriostatic sodium chloride (Hospira, Lake Forest, IL, USA) that contained 52.4 mg/0.2 ml of Cefazolin. At the conclusion of the study, catheters were tested for patency with an infusion of a methohexital/saline solution (5 mg/kg). Rapid loss of muscle tone was interpreted as an indication of patency. Any animal that failed to react to the infusion was excluded from the study.

### Spontaneous Withdrawal

Fentanyl- and vehicle-exposed offspring were tested 24 hours after drug cessation (weaning) on PND 29. Offspring were habituated to the testing room for 30-40 minutes prior to a 15-minute recording session. The session was hand scored by two separate treatment-blinded observers and performed as previously described (de Guglielmo et al., 2014). Recordings were separated into 5-minute bins and withdrawal signs were scored as follows: paw tremors, teeth chattering, genital grooming, ptosis, abnormal gate, excessive/abnormal grooming, trembling, excessive swallowing, and abnormal postures all received 1 point per 5-minute bin. Any behavior that was repeated more than once within a 5-minute bin was given 2 total points. All points from the 3 bins were summed to obtain the total withdrawal score, and the average score between the two observers was calculated to obtain the global withdrawal score.

### Sucrose Splash Test

On PND35 offspring were acclimated to a glass observation cylinder for 5 minutes. Rats were sprayed 3 times (approximately 500 µL total volume) with a 10% sucrose solution on their dorsal coat surface and observed for 5 minutes (Alipio et al., 2021a). The total time spent grooming over the 5-minute period was recorded.

### Von Frey Mechanical Hypersensitivity Test

Mechanical hypersensitivity using the Von Frey apparatus was tested as previously described (de Guglielmo et al., 2019; Simpson et al., 2020). Briefly, on PND40, offspring were acclimated to the testing environment for 10 minutes. Von Frey filaments were applied with increasing force to the hind paws of the animal being tested until the animal sharply withdrew the paw and stopped the measurement. The time of application and total force applied was recorded for each animal across a total of 6 measurements (3 left paw, 3 right paw) and averaged to obtain the paw withdrawal threshold.

### Operant Self-Administration

Offspring were trained to self-administer fentanyl at 8-10 weeks of age. Self-administration (SA) was performed in operant conditioning chambers (29 cm × 24 cm×19.5 cm; Med Associates, St. Albans, VT, USA) that were enclosed in lit, sound-attenuating, ventilated environmental cubicles. The front door and back wall of the chambers were constructed of transparent plastic, and the other walls were opaque metal. Each chamber was equipped with two retractable levers that were located on the front panel. Each session was initiated by the extension of two retractable levers into the chamber. Fentanyl hydrocholoride (3.2 µg/kg per infusion) was delivered through plastic catheter tubing that was connected to an infusion pump. The infusion pump was activated by responses on the right (active) lever that were reinforced on a fixed ratio (FR) 1 schedule, with the delivery of 0.1 mL of the drug per lever press over 4.5 s followed by a 20 s timeout period that was signaled by the illumination of a cue light above the active lever, during which active lever presses did not result in additional infusions. Responses on the left inactive lever were recorded but had no scheduled consequences. Fluid delivery and behavioral data recording was controlled by a computer with the MED-PC IV software installed. Rats were trained to self-administer fentanyl in 12 short access (ShA) sessions (3 hr/day, 5 days/week). At the end of the ShA phase the animals then underwent Progressive Ratio Testing, followed by extinction and reinstatement testing.

### Progressive ratio testing

After FR1 ShA sessions were complete, offspring were tested on a progressive ratio (PR) schedule of reinforcement in which the response requirements for receiving a single reinforcement increased according to the following 1, 2, 4, 6, 9, 12, 15, 20, 25, 32, 40, 50, 62, 77, 95, 118, 145, 178, etc), as previously described (Kallupi et al., 2020). The breakpoint was defined as the last ratio attained by the rat prior to a 60 min period during which a ratio was not completed, which ended the experiment.

### Extinction

After completing 12 days of self-administration and one day of PR testing, offspring were subject to extinction as previously described (Kallupi et al., 2020). In 1-hour long extinction sessions, pressing the previously active lever was no longer associated with any programmed response (no drug infusion and no cue light). The auditory cues were also disabled during extinction sessions. Extinction lasted until responding was at or below 10% of response rates on the final day of ShA self-administration, approximately 8-10 sessions.

### Drug-Induced Reinstatement

Following acquisition of a stable baseline of IV fentanyl self-administration, offpsring were subjected to the extinction phase as described in the prior section (*Extinction*). Immediately following the last extinction session, on the same day, rats were subjected to the drug-induced reinstatement test, as previously described (de Guglielmo et al., 2017; de Wit and Stewart, 1983). All offspring were exposed to a fentanyl priming injection (6.4 µg/kg IV) just before the test session. This test lasted 1 h under conditions identical to those in the extinction phase.

### Cued Reinstatement

Following the drug-induced reinstatement test, offspring repeated the extinction paradigm until responding dropped back to or below 10% of response rates on the final day of ShA self-administration, approximately 5-7 sessions. The following day, rats underwent the cued-reinstatement test (de Guglielmo et al., 2017) in which the conditions were identical to those in the self-administration phase (auditory and visual cues were active) except that fentanyl was not delivered upon active lever responses.

### Corticosterone *ELISA*

Immediately before the final self-administration session, when the rats were presumed to be in withdrawal, blood was collected from the tail vein into tubes containing heparin to measure corticosterone concentrations. Blood samples were spun at 2,500 G for 10 minutes at 4°C to isolate plasma. Plasma was collected and stored at −80°C until analysis. Serum corticosterone (CORT) levels were quantified using the DetectX Corticosterone ELISA kit (Arbor Assays, Ann Arbor, MI, USA). Samples were diluted at 1:100 and plated in duplicate. The assay was completed as described by the kit insert. Average OD values were calculated for each sample, and the mean OD for the NSB was subtracted from each average sample OD value. Sample concentrations interpolated on a 4PL %B/B_0_ standard curve and multiplied by the dilution factor of 100 to obtain neat sample concentrations.

### Brain dissection and RNA extraction

Brain tissues for RNA-seq were collected 1 day after the last IVSA session from 8 prenatal vehicle-treated offspring (6 male, 2 female) and 8 prenatal fentanyl-treated offspring (6 male, 2 female). Animals were euthanized with CO_2_ and immediately decapitated, whole brains were removed from skull and snap frozen. 400 µM thick bilateral CeA brain punches were collected approximately −1.9 mm posterior to Bregma with a 1 mm diameter punch. Tools and cryostat were thoroughly cleaned with EtOH and RNAse Away between samples. Punches were stored frozen at −80°C until further processing. RNA was extracted from the CeA brain punches using Trizol reagent (Invitrogen™) following manufacturer’s instructions. Brain punches were homogenized with Trizol reagent and zirconium oxide beads 0.5 mm RNase Free (Thermo Fisher, NC0284031) using a bullet blender blue (Next Advance, BBX24B) at speed 6 for 2 minutes. After extraction of the RNA, concentrations of RNA were measured using the Qubit Fluorometer (Invitrogen™) and RNA quality was assessed with Agilent 2200 TapeStation and High Sensitivity RNA ScreenTape. RNA samples with RIN above 7 were used for library preparation.

### RNA-seq library preparation and sequencing

NEBNext® Poly(A) mRNA Magnetic Isolation Module (NEB, E7490) and NEBNext UltraExpress® RNA Library Prep Kit (NEB, E3330) were used for RNA-seq library preparation, following manufacturer’s instructions. For each sample, 100 ng of total purified RNA was used as the input amount. A 50-fold diluted adaptor was used for the adaptor ligation step and 12 PCR cycles were performed for the PCR enrichment step. NEBNext® Multiplex Oligos for Illumina® (96 Unique Dual Index Primer Pairs) (NEB, E6448) was used for PCR enrichment reaction to pool each library. Library concentrations were assessed using the Qubit Fluorometer (Invitrogen™). Further, the library quality and sizes of each library were assessed with Agilent 2200 TapeStation and High Sensitivity DNA ScreenTape. Libraries were sequenced on the Illumina NovaSeq X Plus at the IGM Core at UCSD to obtain an average of raw 100bp paired-end reads per sample.

### RNA-seq Bioinformatic Analysis

Raw FASTQ files were subjected to quality control using FastQC.(RRID:SCR_014583). The adaptor sequences and low-quality reads were trimmed using Trimmomatic (Bolger et al., 2014) with LEADING:3 TRAILING:3 SLIDINGWINDOW:4:15 MINLEN:36. Trimmed FASTQ reads were aligned to the rn7 genome (de Jong et al., 2024) using STAR (Dobin et al., 2013) with --outSAMtype BAM Unsorted --outFilterMismatchNmax 5 --outFilterMultimapNmax 1. Reference data were downloaded from UCSC genome browser. After alignment, gene-level read counts were generated using HTSeq (Anders et al., 2015). htseq-count was used for quantification with --stranded=no --idattr=gene_id --type=exon -- mode=union. With quantification data, normalization and differential expressed genes were performed using DESeq2 Gene set enrichment analysis (GSEA) (Subramanian et al., 2005) was performed using the clusterProfiler (Yu et al., 2012) package (v4.14.6) in R (v4.4.2). DEGs were filtered to remove missing values and duplicated gene symbols. A ranked gene list was generated using the log2 fold change values, with gene symbols as names. Gene sets were obtained from the MSigDB database using the msigdbr package (v10.0.0) for the species Rattus norvegicus, including the Hallmark (H), Gene Ontology Biological Process (C5:BP), and Reactome (C2:REACTOME) collections. These sets were combined and converted to a TERM2GENE format required by clusterProfiler. Enrichment was computed using the GSEA() function with the following parameters: a minimum gene set size of 10, a maximum of 500, pvalueCutoff = 0.05, and multiple testing correction via the Benjamini-Hochberg (BH) method. To ensure reproducibility, a random seed was set (set.seed(42)). Pathways with adjusted p-values (FDR) below 0.05 were considered statistically significant.

### Experimental Design & Statistical Analysis

Details regarding the experimental design of individual experiments are provided in the figure legends. All data, with the exception of RNA analysis, were analyzed using Prism 9.0 software (GraphPad, San Diego, CA, USA) and R Studio. Self-administration data were analyzed using one-way ANOVA or two-way ANOVA (time and treatment as factors) with Holm-Sidak’s multiple comparisons *post-hoc* test when appropriate. For pairwise comparisons, data were analyzed using the Student’s t-test. Cohen d effect sizes and 95% confidence intervals were calculated using the R *effsize* package. Missing data due to session failure was imputed as the average from the sessions before and after (<2% of data). The data are expressed as mean ± SEM unless otherwise specified. Values of p < 0.05 were considered statistically significant.

## Results

### Fentanyl-exposed offspring exhibit behavioral and developmental deficits

Perinatal fentanyl exposure significantly impacted pup survival, body weight, and behavior. A survival curve analysis revealed a reduced survival probability for fentanyl-exposed offspring compared to saline controls (Log-rank test, df = 1, χ^2^ = 32.25, p < 0.0001; Fig. 1A). Body weight was assessed at postnatal day (PND) 21, PND35, and PND55, showing a significant time × treatment interaction (two-way repeated measures ANOVA, F(2,112) = 6.394, p = 0.0023) and effect of time (F(1.339,75) = 632.3, p < 0.0001). Holm-Sidak post-hoc tests confirmed lower weights in fentanyl-exposed offspring at PND21 (p = 0.0005) and PND35 (p = 0.0003), but not at PND55 (p = 0.4307) compared to controls (Fig. 1B). Spontaneous withdrawal was evaluated, with fentanyl-exposed offspring displaying higher global withdrawal scores than saline controls (unpaired t-test, t = 15.15, df = 56, p < 0.0001; Fig. 1C). At PND35, negative affect was assessed via the sucrose splash test, revealing reduced grooming time in fentanyl-exposed rats (unpaired t-test, t = 8.039, df = 51, p < 0.0001; Fig. 2A). Mechanical hypersensitivity was tested at PND40 using the Von Frey test, showing a lower paw withdrawal threshold in the fentanyl group (unpaired t-test, t = 2.166, df = 41, p = 0.0362; Fig. 2B). These findings indicate that perinatal fentanyl exposure induces significant developmental and behavioral deficits in offspring.

**Figure 1.**
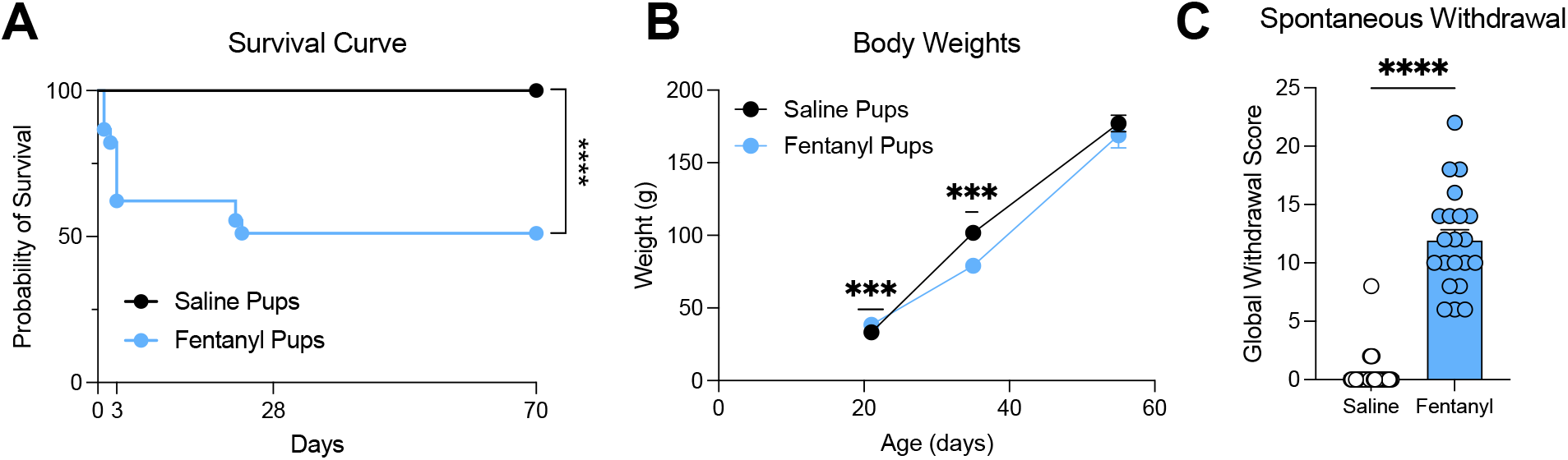
Developmental and withdrawal outcomes in fentanyl-exposed offspring. (A) Survival curves for fentanyl-exposed and saline control offpsring. (B) Body weights at PND21, PND35, and PND55 for both groups. (C) Global withdrawal scores for fentanyl and saline groups. Data represented as mean ± SEM. ***p < 0.001,**** p < 0.0001.

**Figure 2.**
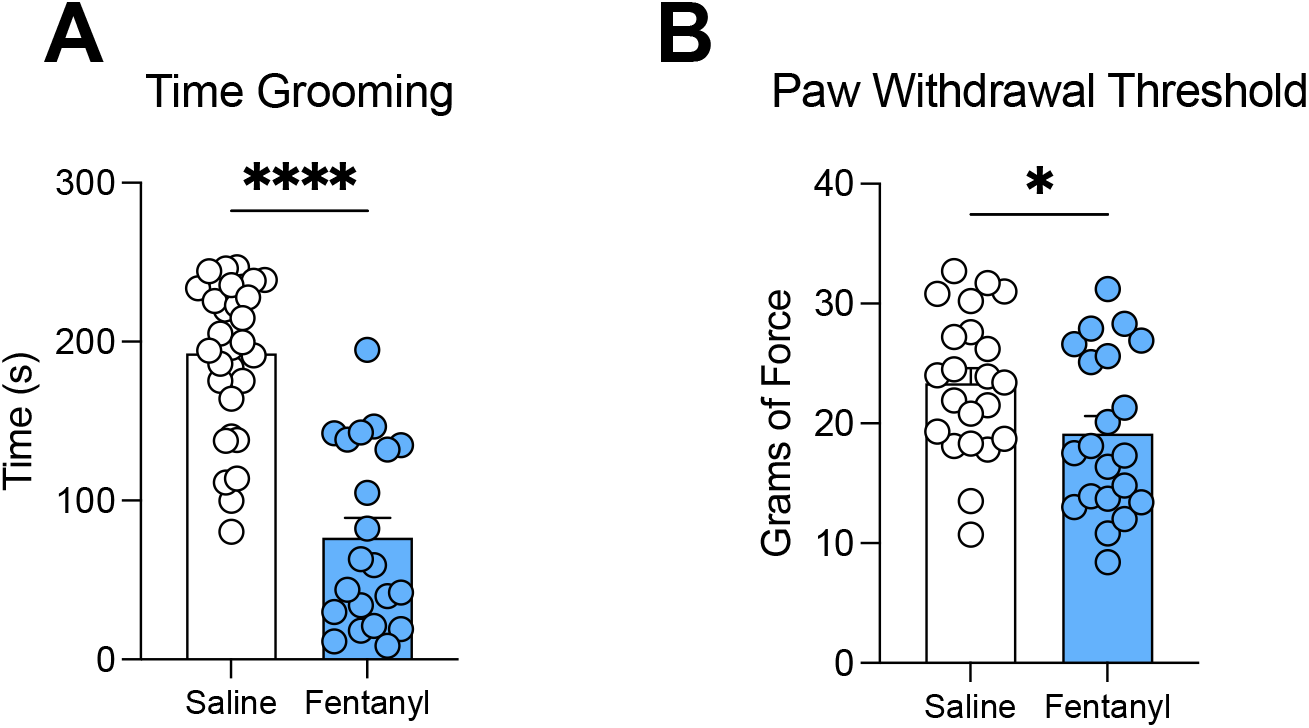
Behavioral outcomes in adolescent offspring. (A) Total grooming time in the sucrose splash test at PND35 for fentanyl-exposed and saline control rats. (B) Paw withdrawal threshold in the Von Frey test at PND40 for both groups. Data represented as mean ± SEM. *p < 0.05,**** p < 0.0001.

### Rats perinatally exposed to fentanyl exhibit disordered drug-taking behaviors as adults

In early adulthood, rats perinatally exposed to fentanyl and vehicle-exposed controls were implanted with jugular catheters and trained to self-administer intravenous fentanyl over 12 days. Fentanyl-exposed rats showed increased fentanyl intake compared to saline controls, with a significant time × treatment interaction (two-way repeated measures ANOVA, F(11,366) = 2.141, p = 0.0171), effect of treatment (F(1,34) = 4.250, p = 0.0470), and effect of time (F(6.111,203.3) = 17.67, p < 0.0001; Fig. 3A). This was not due to non-specific lever pressing, as fentanyl-exposed rats discriminated between reward-associated and inactive levers, showing no significant differences in inactive lever responses (two-way repeated measures ANOVA, no significant effects of time, treatment, or interaction; Fig. 3B). The average number of rewards over the final 3 days of self-administration was higher in fentanyl-exposed rats (unpaired t-test, t = 2.447, df = 34, p = 0.0197; Fig. 3C). No group differences were observed in the progressive ratio test (Fig. 3D). Fentanyl-exposed rats exhibited greater drug-seeking in both drug-induced (unpaired t-test, t = 2.635, df = 34, p = 0.0126; Fig. 3E) and cued reinstatement tests (unpaired t-test, t = 2.662, df = 34, p = 0.0118; Fig. 3F). To quantify plasma corticosterone (CORT) levels during acute withdrawal, a standard curve was established (Fig. 4A; R^2^ = 0.9969). Fentanyl-exposed rats showed elevated CORT levels compared to controls (unpaired t-test, t = 2.255, df = 14, p = 0.0406; Fig. 4B). Correlation analysis revealed a significant positive correlation between CORT levels and fentanyl intake in the fentanyl-exposed group (r^2^ = 0.8723, p = 0.0007) but not in the saline group (r^2^ = 0.3866, p = 0.0998; Fig. 4C).

**Figure 3.**
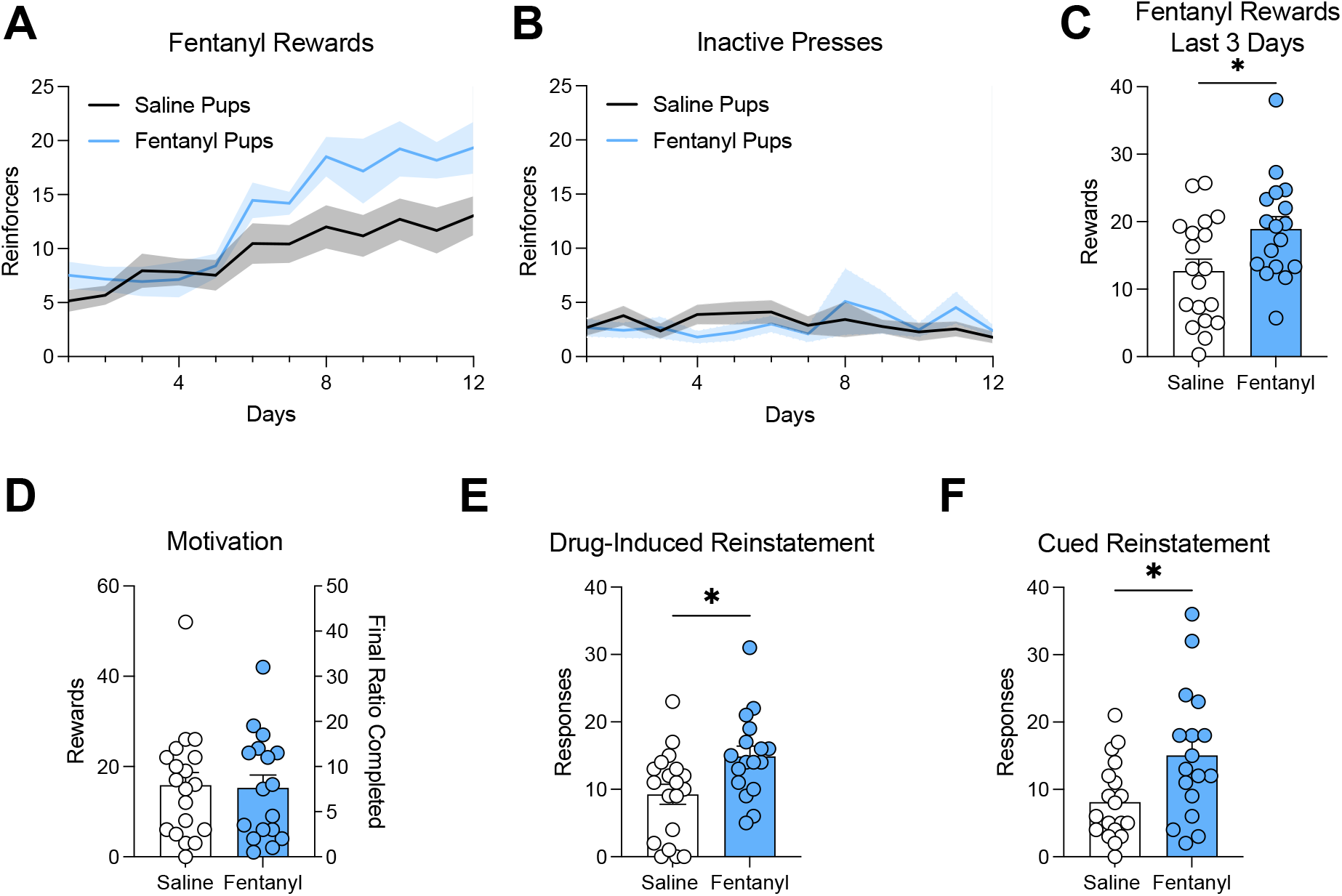
Fentanyl self-administration and reinstatement in adult offspring. (A) Fentanyl rewards earned over 12 days of self-administration for fentanyl-exposed and saline control rats. (B) Inactive lever responses over 12 days. (C) Average number of rewards over the final 3 days of self-administration. (D) Progressive ratio test results. (E) Responses on the drug-associated lever during drug-induced reinstatement. (F) Responses on the drug-associated lever during cued reinstatement. Data represented as mean ± SEM *p < 0.05.

**Figure 4.**
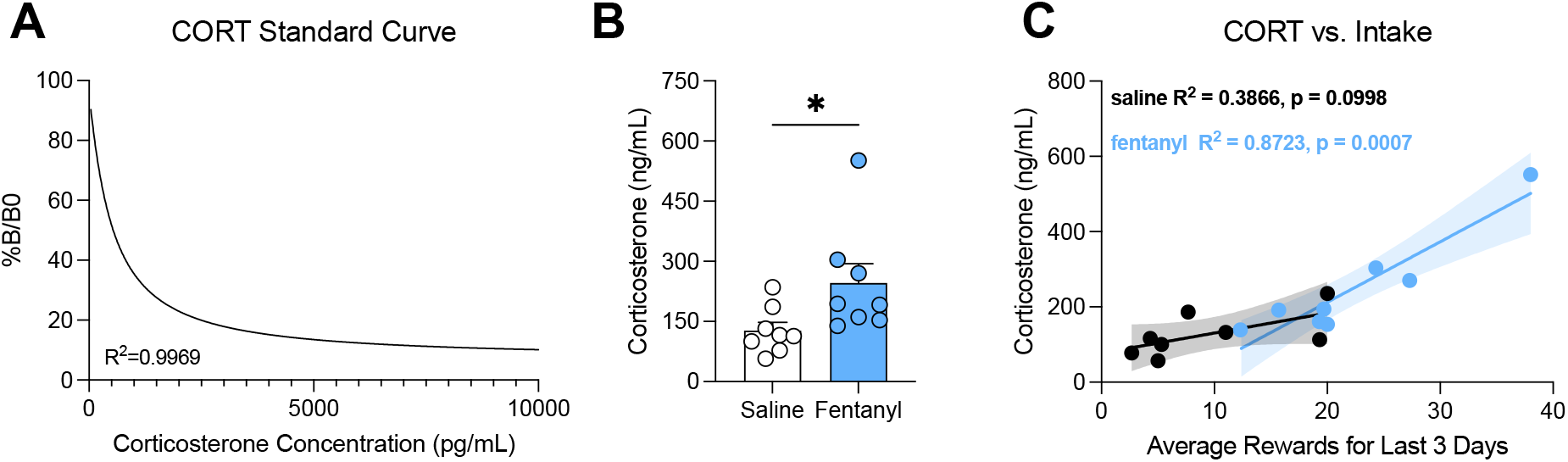
Corticosterone (CORT) levels and correlations in withdrawal. (A) Standard curve for CORT quantification (%B/B0 vs. CORT concentration). (B) Circulating CORT levels in plasma during acute withdrawal for fentanyl-exposed and saline control rats. (C) Correlation between CORT levels and fentanyl intake for fentanyl-exposed and saline groups. Data in (B) represented as mean ± SEM; (A) and (C) show correlation curves.

### Prenatal fentanyl exposure alters gene expression in the CeA

To investigate the long-term transcriptional consequences of prenatal fentanyl exposure, we performed bulk RNA-seq on the CeA of adult rats exposed in utero to fentanyl (n=8, 6 male and 2 female) or vehicle (n=8, 6 male and 2 female) and collected in adulthood 1 day after the end of the fentanyl IVSA protocol. Differential expression analysis identified 110 genes with significant changes in expression (FDR < 0.1, Supplementary Table 1, Fig. 5A). Among these, the top upregulated genes included *Spag6l* (Teves et al., 2014), *Efcab1* (Yamaguchi et al., 2023), and *Ltbp2* (Inoue et al., 2014), while the top downregulated genes included *Rgs22* (Pang et al., 2025). These genes have all been involved in primary cilia functions critical for developmental processes (Youn and Han, 2018). Lastly, several differentially expressed genes (DEGs) were predicted or uncharacterized loci (LOC genes), suggesting the involvement of understudied genomic regions in the brain’s response to perinatal opioid exposure. GSEA revealed 14 pathways significantly enriched (FDR < 0.05, Supplementary Table 2, Fig. 5B) in the CeA transcriptome of prenatal fentanyl-exposed rats. Two gene sets, “voltage-gated potassium channel activity” and “regulation of trans-synaptic signaling”, had positive normalized enrichment scores (NES), indicating upregulation in fentanyl-exposed animals. These pathways are associated with synaptic function and excitability, suggesting potential alterations in CeA neural signaling after perinatal fentanyl exposure. Conversely, the remaining 12 pathways showed negative NES values, reflecting enrichment among genes downregulated by prenatal fentanyl exposure. These downregulated pathways were enriched for developmental processes, including embryonic morphogenesis and anterior-posterior axis. Furthermore, the enrichment of the Class A rhodopsin-like G protein-coupled receptors (GPCR) and GPCR ligand binding gene set with a negative NES suggests that genes involved in GPCR signaling pathways are downregulated in the CeA of rats exposed prenatally to fentanyl. Together, these results indicate that prenatal fentanyl exposure alters gene networks in the CeA of rats, potentially contributing to long-lasting circuit and behavioral alterations in exposed offspring.

**Figure 5.**
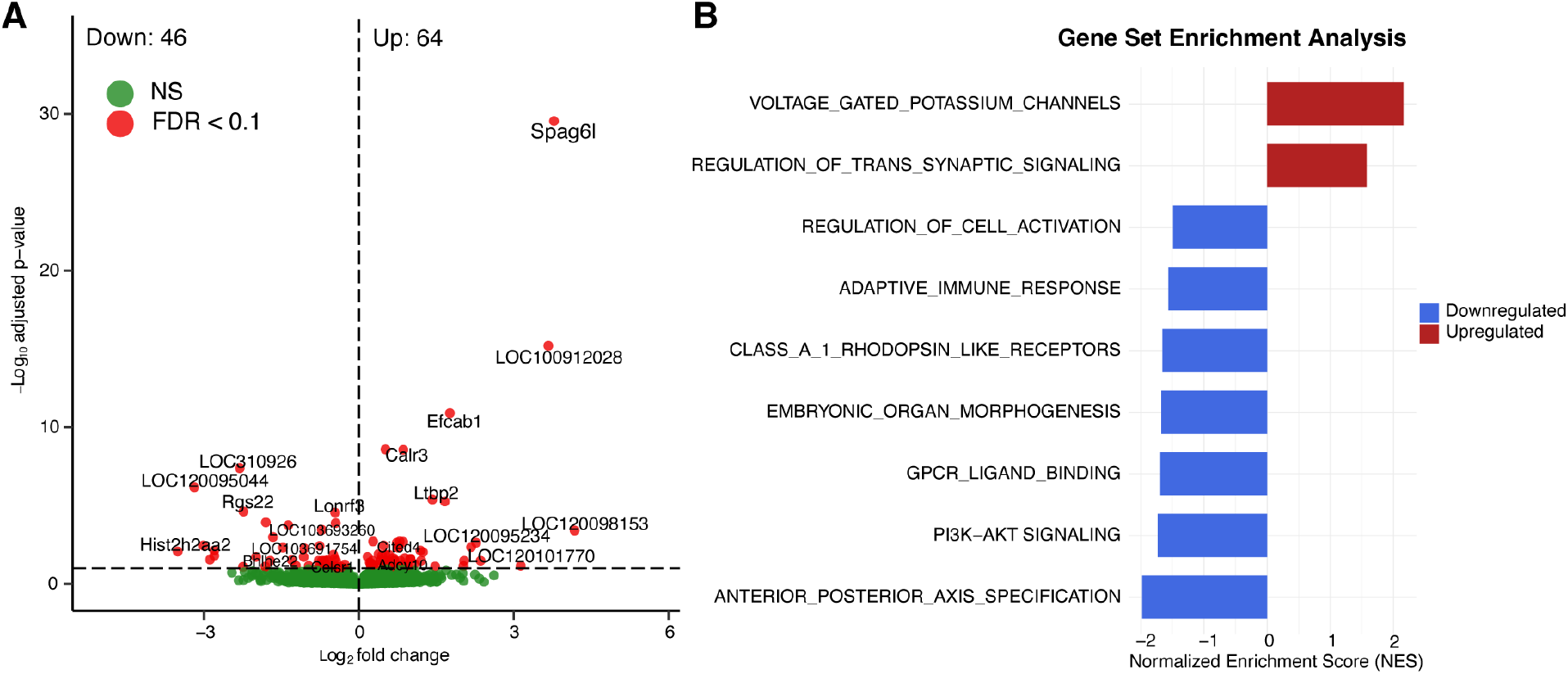
Differential gene expression and pathway enrichment analysis in the amygdala of prenatally fentanyl-exposed rats. (A) Volcano plot showing differentially expressed genes (DEGs) in the central nucleus of the amygdala (CeA) of adult rats exposed to fentanyl in utero compared to vehicle controls. Genes significantly regulated (FDR < 0.1) are highlighted in red, while non-significant genes (NS, FDR ≥ 0.1) are shown in green. (B) Gene set enrichment analysis (GSEA) highlighting significantly enriched pathways (FDR < 0.1). The normalized enrichment score (NES) is plotted, with positive values indicating upregulation and negative values indicating downregulation.

## Discussion

This study elucidates the enduring consequences of perinatal fentanyl exposure in rats, revealing a spectrum of developmental, behavioral, and molecular alterations from birth to adulthood. Fentanyl-exposed offspring exhibited reduced survival, lower body weight, and heightened withdrawal symptoms, followed by adolescent negative affect and mechanical hypersensitivity, and adult increases in fentanyl self-administration, drug-seeking, and CORT levels. RNA-seq analysis of the CeA identified dysregulated genes and pathways, offering mechanistic insights. These findings advance prenatal opioid research in rodents by tracking litters into adulthood, elucidating the long-term consequences of fentanyl exposure.

Neonatal outcomes in our model, such as reduced survival, lower body weight, and elevated withdrawal scores, mirror NOWS symptoms in human infants exposed to opioids in utero, consistent with clinical reports (Ko et al., 2016). Rodent models of prenatal morphine or buprenorphine exposure similarly show growth deficits and withdrawal behaviors, validating our model’s relevance (Myers et al., 2024; Searles et al., 2025; Wallin et al., 2019). Unlike most studies focusing on early development, our work extends to later stages, demonstrating fentanyl’s enduring effects on behavior and neural gene expression.

In adolescence, fentanyl-exposed rats displayed negative affect (reduced grooming in the sucrose splash test) and mechanical hypersensitivity (lower paw withdrawal thresholds), indicating disrupted emotional and sensory processing. These findings align with clinical observations of internalizing behaviors and altered pain perception in opioid-exposed children (Conradt et al., 2019). Preclinical studies report similar sensory deficits in fentanyl-exposed mice, suggesting conserved effects across opioids (Alipio et al., 2021b).

Adult rats in our study exhibited heightened addiction vulnerability, with increased fentanyl self-administration, enhanced drug-seeking during reinstatement, and elevated CORT levels, correlating with intake. These results suggest altered reward and stress circuits, supported by preclinical evidence of enhanced drug self-administration following prenatal morphine exposure (Glick et al., 1977; Ramsey et al., 1993). Human studies also indicate increased neuropsychiatric risks in opioid-exposed children, reinforcing our findings (Kang et al., 2024). By examining fentanyl self-administration, our study establishes a direct link between prenatal exposure and increased addiction-like behaviors in adulthood with the same opioid.

To elucidate the molecular underpinnings of heightened addiction vulnerability and elevated CORT levels observed in adulthood, we performed RNA-seq analysis on the CeA, a critical brain region modulating stress responses and reward-seeking behaviors (Koob, 2009) implicated in opioid dependence (Chaudun et al., 2024; Kallupi et al., 2020). The observed transcriptional dysregulation of genes associated with neurodevelopmental processes, coupled with the significant downregulation of GPCR ligand binding pathways in the adult CeA of the amygdala following perinatal fentanyl exposure, suggests that disrupted neurodevelopmental processes and GPCR signaling mechanisms may contribute to the behavioral alterations observed in this rat model. These transcriptomic changes align with findings in opioid-exposed rodents, where neonatal morphine exposure alters synaptic and developmental gene networks in the forebrain, affecting circuit maturation (Dunn et al., 2023).

Similarly, perinatal fentanyl exposure in mice induces molecular changes in the somatosensory cortex (Alipio et al., 2021b), disrupting sensory processing and supporting the notion of widespread neural impacts across brain regions. Moreover, the GSEA revealed reduced adaptive immune response in the CeA of rats exposed to fentanyl prenatally. These effects could be a consequence of the elevated CORT levels associated with prenatal opioid exposure, which are known to suppress immune functions (Xu et al., 2025).

Among the most notable gene expression changes was the downregulation of *Rgs22* and upregulation of *Gpr88. Rgs22* is a member of the regulator of G protein signaling (RGS) family, which regulates ependymal cell function to prevent hydrocephalus in mice (Pang et al., 2025), suggesting an important developmental role. RGS proteins function as GTPase-activating proteins (GAPs) that attenuate GPCR signaling, a process critical for neuromodulation. Interestingly, RGS2, a close paralog of *Rgs22*, has showed genetic associations with anxiety, panic disorder, and suicidality in early humans studies (Cui et al., 2008; Hohoff et al., 2015; Mouri et al., 2010). *Gpr88* is a striatum-enriched GPCR that regulates motor function, mood, and reward learning (Ye et al., 2019). Recent studies suggest it modulates opioid-related behaviors, including morphine sensitivity and reward processing, highlighting its potential relevance to addiction biology (Laboute et al., 2020; Ye et al., 2019). Human genetic studies have suggested that *RGS22* is associated with educational attainment (Okbay et al., 2022).

A key limitation of this study is that all animals, regardless of prenatal treatment, were subsequently exposed to fentanyl through fentanyl IVSA in adulthood. Notably, rats exposed to fentanyl in utero exhibited higher levels of fentanyl intake during the IVSA phase compared to controls, raising the possibility that altered drug-taking behavior in adulthood may have amplified or confounded the transcriptional effects attributed to prenatal exposure. Future studies using drug-naive adult animals or a yoked control design will be necessary to isolate the specific contributions of prenatal opioid exposure from the cumulative effects of postnatal drug intake.

In summary, this study demonstrates that perinatal fentanyl exposure in rats induces persistent developmental, behavioral, and molecular alterations, with significant implications for understanding NOWS and addiction vulnerability. The observed neonatal deficits, adolescent emotional and sensory dysregulation, adult addiction-like behaviors, and CeA transcriptomic changes underscore the profound impact of prenatal fentanyl on brain development and function. Unlike human studies, prenatal fentanyl exposure was randomly assigned, and all other aspects of the environment were similar for fentanyl exposed and control offspring. These findings provide a robust preclinical framework for investigating the long-term consequences of synthetic opioid exposure, supporting the need for targeted interventions to mitigate NOWS and associated neuropsychiatric risks in human populations exposed to fentanyl during pregnancy.

## Abbreviations

CeA: Central Nucleus of the Amygdala
CORT: Corticosterone
DEG: Differentially Expressed Genes
FR: Fixed Ratio
GPCR: G-Protein Coupled Receptor
HS: Heterogeneous Stock
IV: Intravenous
INSA: Intravenous Self Administration
NOWS: Neonatal Opioid Withdrawal Syndrome
PR: Progressive Ratio
ShA: Short Access

## Data Availability Statement

All data that support the findings of this study are available upon request.

## Author Contributions

CPW conceptualized and performed and oversaw all in vivo experiments, performed data analysis, and wrote the manuscript. MB, PC, and SD assisted with all in vivo experiments and surgical procedures. BM, GC, GP, PN, AM, and SRMUR assisted with animal handling and data collection. FT and YS performed RNAseq and analysis and contributed to the manuscript. AP provided HS rats and contributed to the manuscript. GdG assisted with experimental design and contributed to the manuscript.

## Funding

Research is funded by the National Institutes of Health (R01DA056602 and P30DA060810).

## Competing Interests

The authors have nothing to disclose.

## Notes

### Competing Interest Statement

The authors have declared no competing interest.

### Summary of Updates

one name was mispelled and one funding source was missing

